# Neural computations combine low- and high-order motion cues similarly, in dragonfly and monkey

**DOI:** 10.1101/240101

**Authors:** Eyal I. Nitzany, Gil Menda, Paul S. Shamble, James R. Golden, Qin Hu, Ron R. Hoy, Jonathan D. Victor

## Abstract

Visual motion analysis is fundamental to survival across the animal kingdom. In insects, our understanding of the underlying computations has centered on the Hassenstein-Reichardt motion detector, which computes two-point cross-correlation via multiplication; in mammalian cortex, it is postulated that a similar signal is computed by comparing matched squaring operations. Both of these operations are difficult to implement biophysically in a precise fashion; moreover, they fail to detect the more complex multipoint local motion cues present in the visual environment. Here, via single-unit recordings in two visual specialists, dragonfly "(Odonata)" and macaque, and via model simulations, we show that neuronal computations are not simply approximations to idealized behaviors forced by biological constraints, but rather, are signatures of a common computational strategy to capture multiple local motion cues. The similarity of motion computations at the neuronal level in the brains of two extremely dissimilar animals, with evolutionary divergence of over 700 Myr^1^, suggests convergence on a common computational scheme for detecting visual motion.

## Introduction

Across the animal kingdom, analysis of motion in the visual input is crucial for survival. Because of its life-criticality and prevalence, this process has become an important model system for understanding how algorithms are implemented in neural circuitry.

Motion analysis is generally thought to begin with a stage in which local motion signals are extracted. Especially in insects, our understanding of this first stage has centered on the notion of an “elementary motion detector” (EMD), a circuit that extracts these signals by correlating the visual input at one point, with the visual input at a second point and a later time^2^. Faithful implementation of this two-point cross-correlation corresponds to multiplication of neural signals. However, detailed analysis of neural circuitry in the fly^3,4^ indicates that multiplication of neural signals is only approximate.

This begs the question of whether deviations from the ideal multiplication central to the EMD constitute a limitation of neural hardware, or rather, expressions of computational subtlety that have evolved to exploit the characteristics of the visual environment^5^. Interestingly, recent work points to the latter. Fitzgerald and Clark^6^ showed that self-motion in a naturalistic environment generates optic flow signals that would not be detected by an EMD, along with the expected motion signals that an EMD is built to detect. These authors further showed that motion detectors with specific deviations from multiplication could capture the signals that the EMD would miss. Such motion detectors were more accurate in extracting motion flow, and more predictive of the optomotor behavior of flies, than a standard (i.e., precisely multiplicative) EMD.

While these findings suggest that deviations of neural computations from strict multiplication are functionally important, it is unclear where the results of these computations emerge, and how they are executed. For example, the behavioral data leaves open the possibility that the retinal output has the properties expected of an EMD, and the additional motion signals needed to account for behavior arise elsewhere. Moreover, it is unclear whether the basic conclusion that these deviations are a “feature” rather than a “bug” would extend from fruit flies to organisms that are visual specialists – that is, organisms that rely on motion signals for the demanding tasks of identifying and capturing prey, and not simply for the generic tasks of extraction of optic flow and avoidance of predators.

Here, we address these questions by showing that different kinds of motion signals coexist in individual neurons in early visual areas of the brains of two widely divergent visual specialist species: the medulla and lobula of the dragonfly, and the V1 and V2 of the macaque monkey, animals separated by ~700 Myr of evolutionary history^1^. These species have profound differences in eye morphology, brain size, and brain organization. Moreover, these species are separated by a fundamental difference in the way that directionally-selective motion signals arise: in the dragonfly^7^ as in drosophila^8^, via a multiplication-like operation at the retinal level; in the macaque^9,10^, via subtraction of matched squaring operations in visual cortex. Despite these differences, we show that the pattern of neural sensitivity to local motion signals have a number of similarities. Identification of these parallels in such widely divergent visual specialist species suggests that the algorithms used for local motion extraction by neural circuits reflect a convergent evolutionary process.

Perhaps counterintuitively, characterizing the neural computations underlying motion detection is facilitated by the use of motion stimuli that are uncommon in nature. The logic is that while different species may use diverse algorithms, stimuli that are common in the environment are expected to produce near-veridical outputs. Thus, in order to distinguish among the internal workings of successful algorithms, it is helpful to use stimuli that are “out of sample” – i.e., un-natural stimuli, on which the algorithms have not been trained, and for which the algorithms may fail in different ways. Along with several other recent studies^3,5,6^, we use a mathematical dissection of the potential cues to visual motion (Figure 1) to create a battery of such stimuli. We describe the strategy briefly here; further details are in ^5,11,12^. The starting point is the intuition behind the EMD: that a correlation between one point and a second point, offset in space and time, is a cue to visual motion. However, correlations between any number of points in a slanted spatiotemporal region can also indicate motion. This was emphasized by Chubb and Sperling^13^, who constructed stimuli that drive a visual motion percept based on correlations among four points. They called this phenomenon “second-order” or “non-Fourier” (NF) motion. (The term NF motion arose to distinguish these signals from motion signals that can be detected by the standard EMD; the latter is also called Fourier (F) motion, as it can be extracted from the amplitude of the stimulus’s Fourier components.) Subsequently, stimuli that drive percepts of motion based on correlations among three points were identified by Hu and Victor^11^, a phenomenon that has become known as “glider” (G) motion. G motion is also undetectable from the Fourier amplitudes and hence by a standard EMD, but we give it a separate designation because it is also undetectable by the mechanisms postulated by Chubb and Sperling^13^. These several kinds of motion signals and their subtypes typically co-occur in nature^12^, but it is possible to generate un-natural, artificial stimuli that isolate each of them, thus allowing measurement of responses to each kind of motion signal, and providing a “fingerprint” of the motion computations. These artificial stimuli form the basis of the present study.

**Fig. 1.**
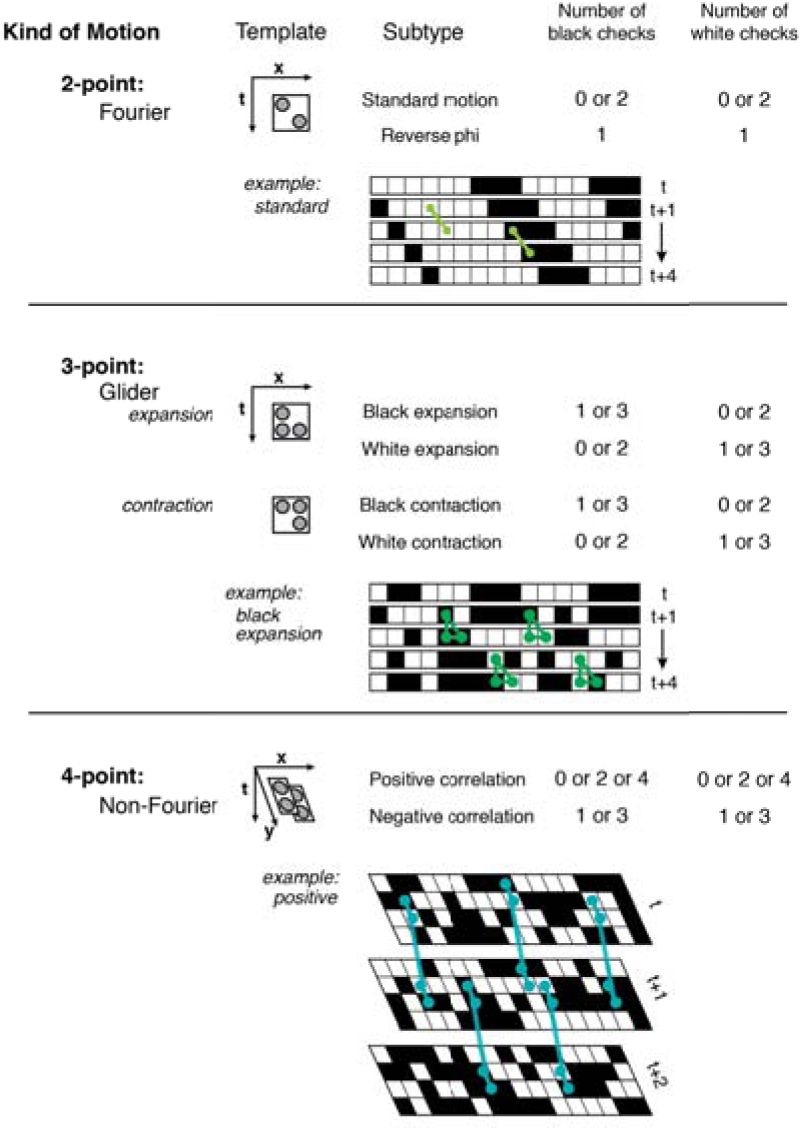
Summary of motion stimuli. Each kind of motion (first column) corresponds to a “template” (second column) containing two, three, or four points in space and time. For each kind of motion, there are two or four subtypes (third column), which are defined by parity rules (last two columns), and, for glider motion, the orientation of the template. One subtype is illustrated in detail for each kind of motion; movies of all subtypes can be found in the video supplement. ***Fourier* (*F*) *motion*** (**top panel**): the template consists of two checks on a space-time diagonal. In the illustrated “standard motion” subtype, there are always 0 or 2 black checks within the template, so check colors match along the diagonal. In the “reverse-phi” subtype, the number of black checks within the template is always 1, so check colors alternate along a diagonal. ***Glider* (*G*) *motion*** (**middle panel**): the template contains three checks in a space-time triangle. For “expansion” subtypes, the triangle is oriented so that it expands as time progresses; for “contraction” subtypes, the template orientation is reversed. In the illustrated “black expansion” subtype, there are always 1 or 3 black checks within the template. This generates a stimulus containing expanding black regions. ***Non-Fourier* (*NF*) *motion*** (**bottom panel**): the template contains four checks in a space-time parallelogram. In the illustrated “positive correlation” subtype, the number of black checks within the template is always 0, 2, or 4. This generates a stimulus in which an edge between the two checks in the template at time *t* is always followed by an edge between the other two checks of the template at time *t*+1. In the “negative correlation” subtype, the number of black checks in the template must be 1 or 3. This generates a stimulus in which edges within the template are present either at time *t* or *t*+1, but not both.

While the initial motivation for the approach is mathematical, two considerations point to the functional significance of these several motion types. First, G motion signals are generated when objects are looming or receding. Second, these three kinds of motion signals (F, G, and NF) are known to elicit behavioral responses in a wide range of species (*Drosophila*:^3^; zebrafish:^14^; human:^11,15^); for two of the kinds of signals (F and NF), neurophysiologic correlates in the mammalian visual cortex are documented (macaque:^16^; cat:^17^).

Crucially, different candidate models for motion computations make sharply contrasting predictions for this stimulus set – as previously recognized ^6,11,18^, and as we further demonstrate here with an expanded set of models. For example, a strict Reichardt EMD model does not respond to G stimuli, but modifying its nonlinearity can lead to models that respond selectively to NF or G stimuli, and with different sensitivities. In sum, probing a neural system with a battery of movies that isolate different types of motion signals – including those detectable by a standard EMD and those that are not – is an effective way to characterize its computations.

We deployed this approach while recording the activity of individual spiking neurons in visual brain areas of the dragonfy (medulla and lobula) and the macaque (cortical areas V1 and V2). For recording in the dragonfly, we used a recently-developed technique^19^ to enable efficient extracellular recording with sharp metal electrodes; in the macaque, we used a standard tetrode technique^20^. We then used standard spike sorting and data analysis procedures to quantify the sensitivity of individual neurons in each brain area to the several kinds of motion stimuli. What emerged is that despite the obviously vast morphological differences in their eyes (compound vs. simple lensed), brain size, brain organization, number of neurons, body plan, life histories, and over 700 Myr of evolutionary separation^1^, dragonfly and macaques have neurons that are remarkably similar in their responses to multiple kinds of motion cues. While there are some differences in the motion fingerprints between species, these differences are minor in comparison with the variety of fingerprints that are encountered in the space of models that generalize the EMD. These parallels suggest that the specific kinds of deviations encountered in these two divergent species result from a convergent adaptation to statistical features of motion in the natural environment.

## Results

To analyze motion processing in the brains of the dragonfly and macaque, we probed their visual systems with representatives of the diverse motion signals that exist in natural stimuli ^5,12^ For recording, we used metal microelectrodes to record visual responses from the medulla or lobula complex of the dragonfly and primary and secondary visual cortices (area V1 or V2) of the monkey. These extracellular recordings can then be parsed, separating single units on the basis of their amplitude and waveform using spike-sorting software (see Methods).

Example visual unit responses are shown in Figures 2 and 3. The first example from the dragonfly medulla (Figure 2, left, top) has a directionally-selective response to standard Fourier motion, but not to any of the other motion stimuli. However, this pattern of behavior occurred only in a minority of the directionally-selective neurons. More commonly, neurons were responsive to multiple kinds of motion signals. For example, the second medulla neuron in Figure 2 (left, bottom), was selective for standard motion in one direction, and black glider contraction in the opposite direction. The example neurons from the dragonfly lobula (Figure 2, right) are further instances of neurons that are sensitive to multiple motion types. Both have directionally-selective responses to standard F motion and directionally-selective responses in the opposite direction to reverse-phi F motion. Both lobula neurons also have directionally-selective responses to G expansion, and the preferred direction for G expansion depends on the polarity (black vs. white) of the glider: it is the same as for F motion for a black expanding glider stimulus, and opposite for a white one. In addition, the second lobula neuron (right, bottom) also has a small but statistically significant directionally selective response to black G contraction.

**Fig 2.**
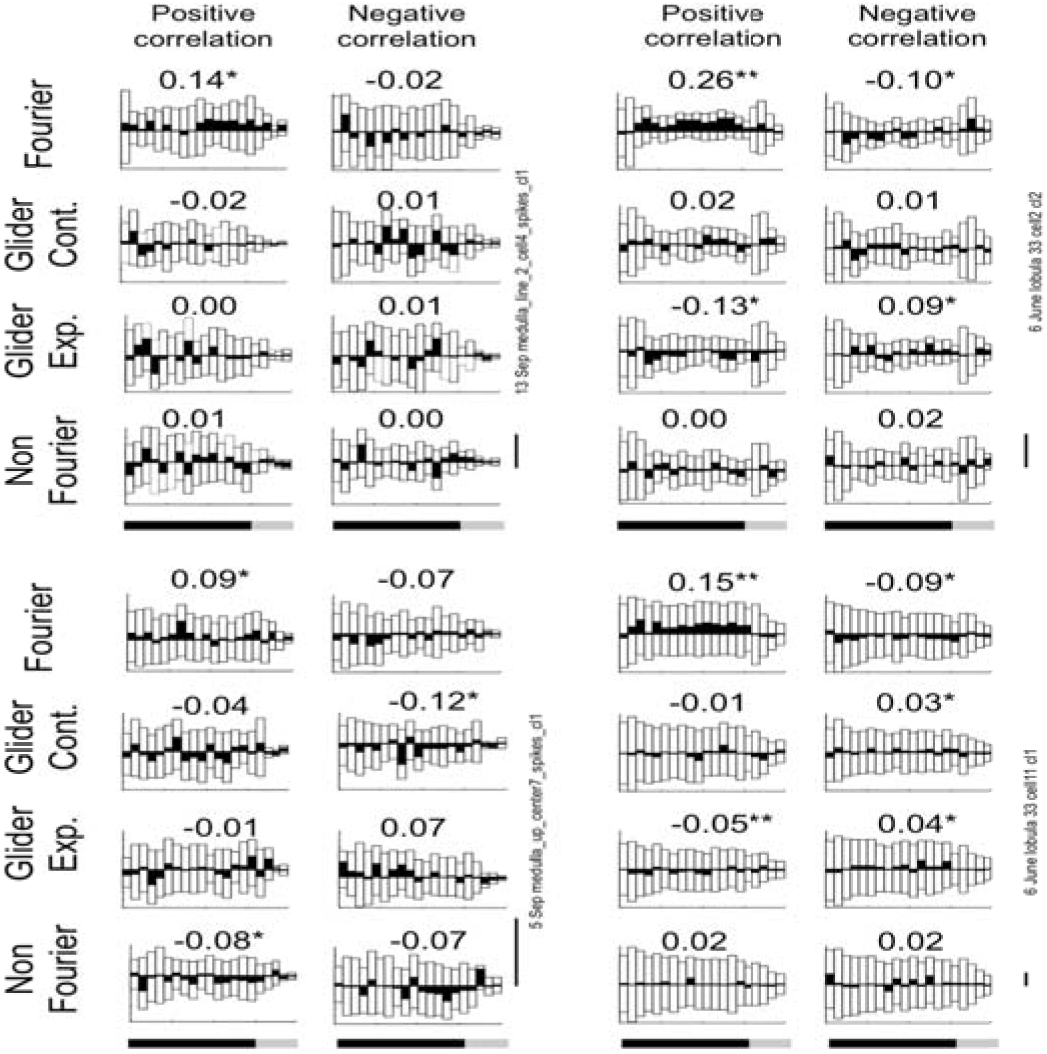
Responses of example neurons in dragonfly medulla (left) and lobula (right). Each set of eight plots shows responses of one neuron to several kinds of motion stimuli: the four rows correspond to the type of motion (Fourier (F), glider (G) contraction, G expansion and non-Fourier (NF)), and the two columns correspond to the correlation sign (see Figure 1). For F motion, positive correlation is standard motion, negative is reverse-phi. For G motion, positive correlation has white spatiotemporal triangles, negative correlation has black spatiotemporal triangles. For NF motion, positive correlation means an “even” constraint on four luminance values; negative correlation means an “odd” constraint. For each of these 8 stimulus types, 32 example movies were presented in the direction preferred for F motion (top), and in the opposite direction (underneath). Presentations lasted 1500 ms and were followed by 500 ms of a gray background. Open histograms show the responses in each direction, the black histogram shows the difference. The DI, indicated for each stimulus type, is computed in the standard fashion (*M_pref_* – *M_non_._plef_*)*/* (*M_prsf_* + *M_non_._pref_*). For each motion type, the sign of the direction selectivity index (DI) indicates the preferred direction relative to the preferred direction for Fourier motion: positive if it is the same direction, negative if it is opposite. Significance (via paired t-test) is indicated by asterisks: *: p<0.05, **, p<0.01. Scale bar is 20 impulses/sec.

**Figure 3.**
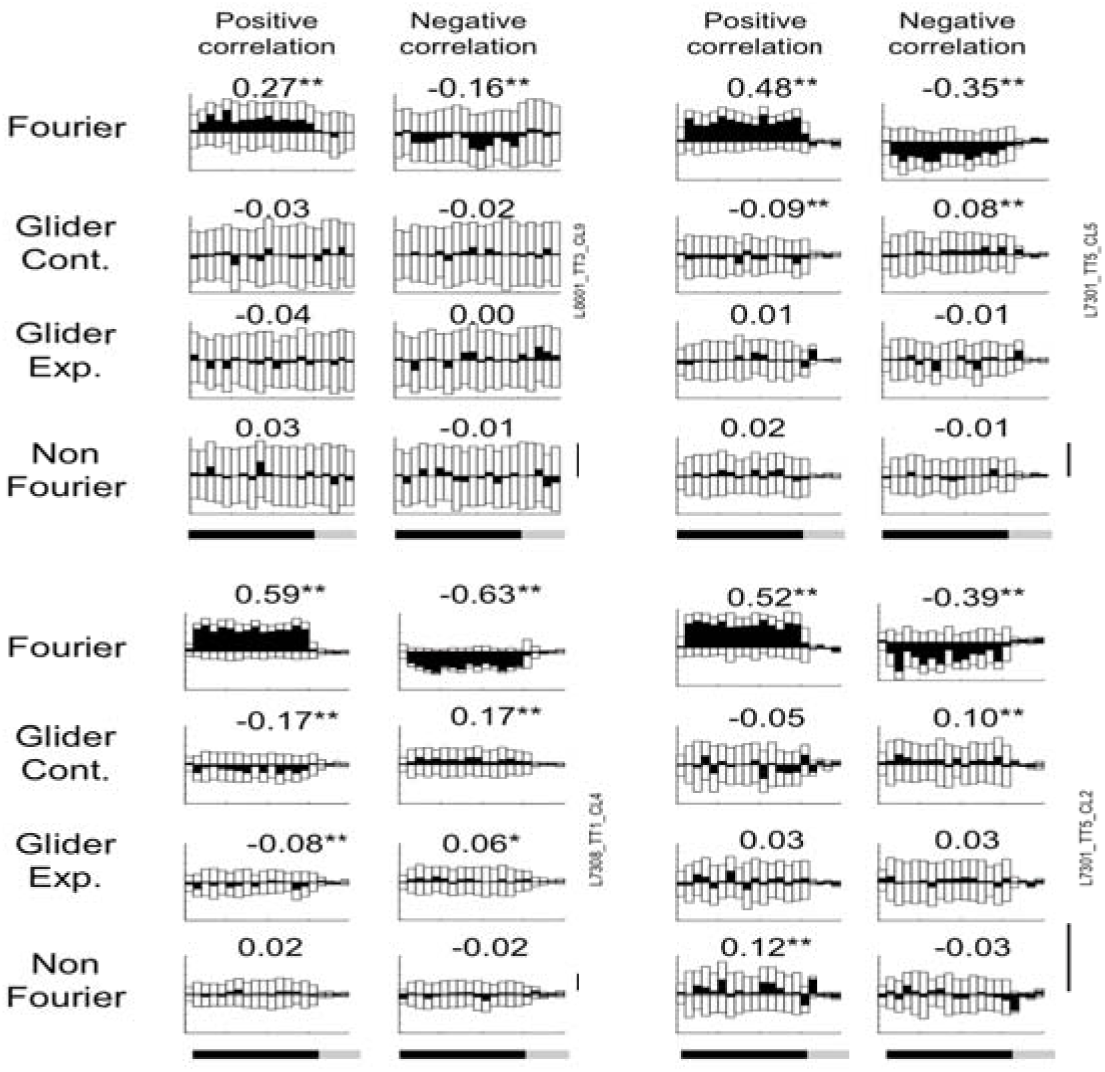
Responses of example neurons in macaque V1 (left) V2 (right). Scale bar is 10 impulses/sec. Stimulus details and other plotting conventions as in Figure 2.

Here and below, we quantified motion selectivity two ways (see Methods for further details): by a paired t-test comparing responses to motion stimuli in opposite directions (see Methods for further details), and by the standard Direction Selectivity Index (DI)^21^, defined by

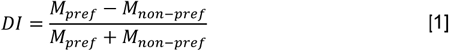
where *M_pref_* is the firing rate elicited by stimulus movement in the direction that is preferred for F motion, and *M_non_._pref_* is the firing rate for stimulus movement in the opposite direction. Thus, *DI*=0 indicates a cell has no direction selectivity and a *DI*>0 indicates the presence of direction selectivity for F motion (with maximum value of 1). We define the DI similarly for the other kinds of motion, but maintain the convention that *M_pref_* is the response in the direction preferred for F motion. Thus, *DI>0* indicates a motion preference in the same direction as for F motion and *DI*<0 indicates a motion preference in the opposite direction. For example, for both neurons in the right half of Figure 2, the DI for black G expansion is positive, while the DI for white G expansion is negative, indicating that the cells’ direction preference for the former is the same as for F motion, and the cells’ direction preference for the latter is opposite. (Since we fix the “preferred direction” according to the responses to the F stimulus, the DI for F motion must be non-negative.)

As in the dragonfly recordings, the macaque recordings revealed neurons that were sensitive just to F motion, as well as neurons that were sensitive to multiple motion types. For example, the first V1 illustrated neuron (Figure 3, left, top) responds to standard F motion and to reverse-phi motion (but with opposite direction selectivity), while the second V1 neuron (Figure 3, left, bottom) additionally responds to all subtypes of G motion. The direction selectivity depends on the polarity (white vs. black) of the glider, but not on whether it is expanding or contracting. The first V2 example neuron (Figure 3, right, top) responds to standard F motion and G contraction – again, with opposing direction selectivities, depending on polarity. The second V2 example (figure 3, right, bottom) responds to standard and reverse-phi F motion, and in addition has a comparatively weaker directionally selective response to NF motion and black G contraction. The latter two responses both have the same direction preference as standard F motion.

### Patterns of responses to motion subtypes in dragonfly and macaque

As an initial step to summarize and compare the patterns of motion sensitivity encountered in these neural populations, we determined the fraction of neurons with directionally-selective responses to each kind of motion. Here, we considered a neuron to have a directionally selective response to a motion type (F, G, or NF) if there was a statistically significant difference between the firing rates elicited by opposite motion directions for any of its subtypes (e.g., standard or reverse phi for F, any of the four subtypes for G, and either positive or negative correlation for NF – see Figure 1). Figure 4 summarizes the prevalence of neurons with directionally-selective responses to F and G motion stimuli, as well as to analogous non-Fourier (NF) motion stimuli. As exemplified by Figures 2 and 3, each kind of stimulus elicits directionally-selective responses in the medulla and lobula of the dragonfly, and in V1 and V2 of the macaque. The summary of Figure 4 shows that in the two species, the proportions of neurons that are sensitive to each motion type are similar.

**Figure 4.**
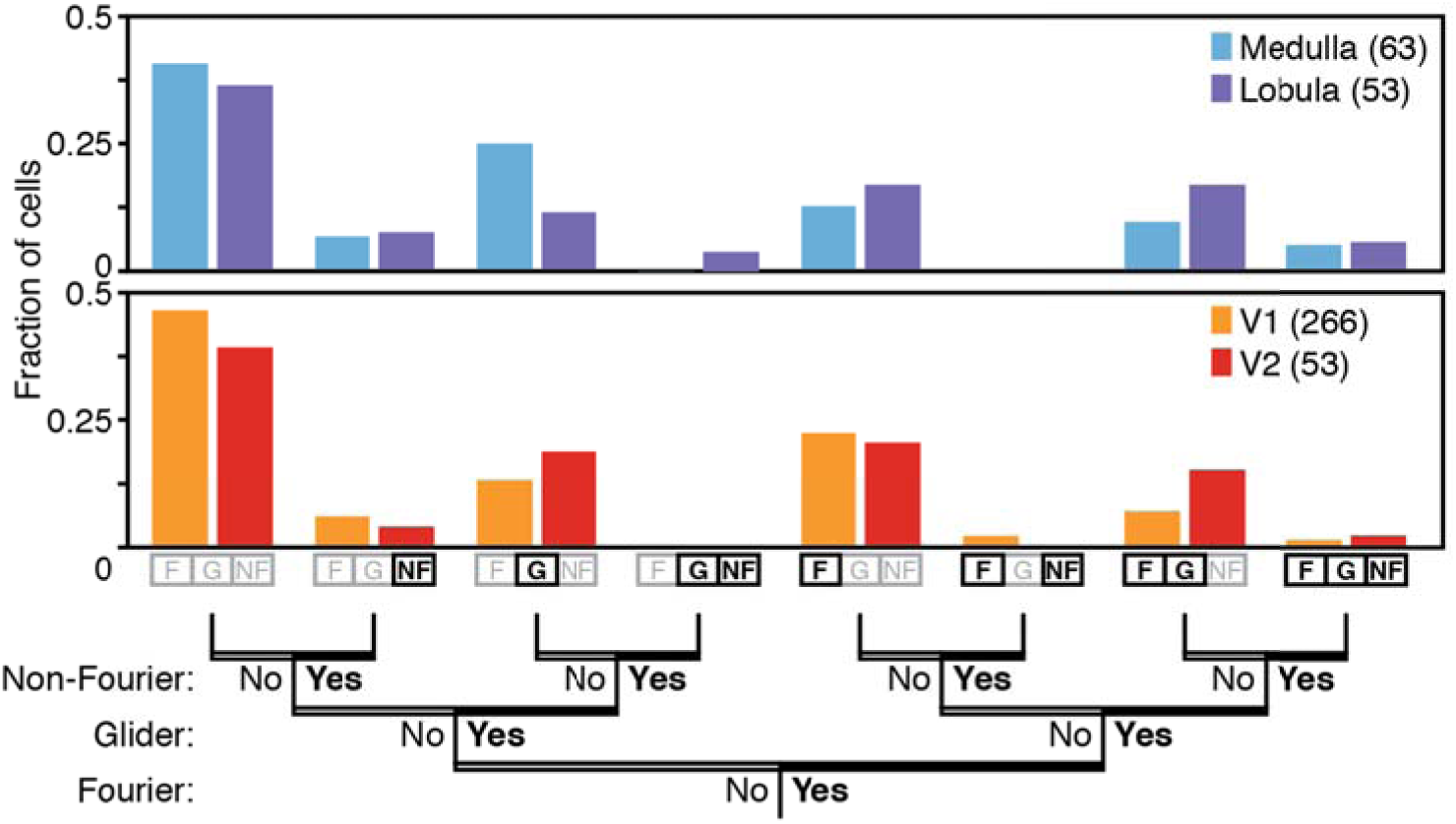
Similar fractions of central neurons respond to Fourier, glider, and non-Fourier motion signals in dragonfly and macaque. Neurons were considered to have a significant response to a kind of motion if their firing rate was significantly larger for stimuli with motion in one direction than in the other (p<0.05, two-tailed paired t-test). In both species and for both brain regions, similar fractions of neurons responded to each of the three major motion signals, and similar fractions of neurons responded to more than one kind of motion. In both species, many cells did not respond selectively to any motion stimulus (far left bars), as is typical of recordings from the visual brain. See Supplementary Figure 1 for analysis using an alternative criterion for sensitivity.

This analysis also shows that the neural circuitry for analyzing local motion signals does not follow the mathematical partitioning used to construct the stimuli. While the segregation between F, G, and NF motion signals is a mathematically clean one, the recordings show that they are jointly analyzed: many neurons respond to more than one motion type. Notably, the frequency of neurons that respond to each combination of motion types is similar across species. Figure S1 shows that the same conclusion is reached using a different statistical criterion for sensitivity to each motion type, in which a multiple-comparisons correction is made within each motion type.

We next consider a more detailed analysis of the neural populations, focusing on the degree and direction of sensitivity to the multiple motion types within individual neurons. Figure 5 shows how each neuron’s sensitivity to the several motion types relates to their responses to standard F motion. In these plots, the sign of the DI for the nonstandard motion types indicates whether the preferred direction was the same as (DI >0) or opposite to (DI <0) the preferred direction for standard F motion. In both species, neurons that have directionally-selective responses for one subtype often have directionally-selective responses for another, but the direction may differ. Specifically, neurons that responded to standard F motion in one direction tended to respond to black G contraction in the same direction, but white G contraction in the opposite direction. The same correspondence held for black and white G expansion, but the correlation for macaque responses was weaker. Both species also showed a strong anticorrelation between selectivity to standard F motion and reverse-phi motion. For NF stimuli, though, the two species behaved differently: F sensitivity and NF sensitivity were positively correlated in macaque, but negatively correlated in dragonfly. All of these correlations were significant (p<0.05, via t-statistics, computed by Matlab’s regstats), except for white G contraction in the dragonfly and black G expansion in the macaque. Notably, the neuronal pattern of motion sensitivities corresponds to findings at the behavioral level in the fruitfly^6,18^: Fitzgerald, Clark, and colleagues found that fly optomotor responses to black G expansion and contraction were in the same direction as responses to F motion, while responses to white G expansion and contraction were in the opposite direction.

**Figure 5.**
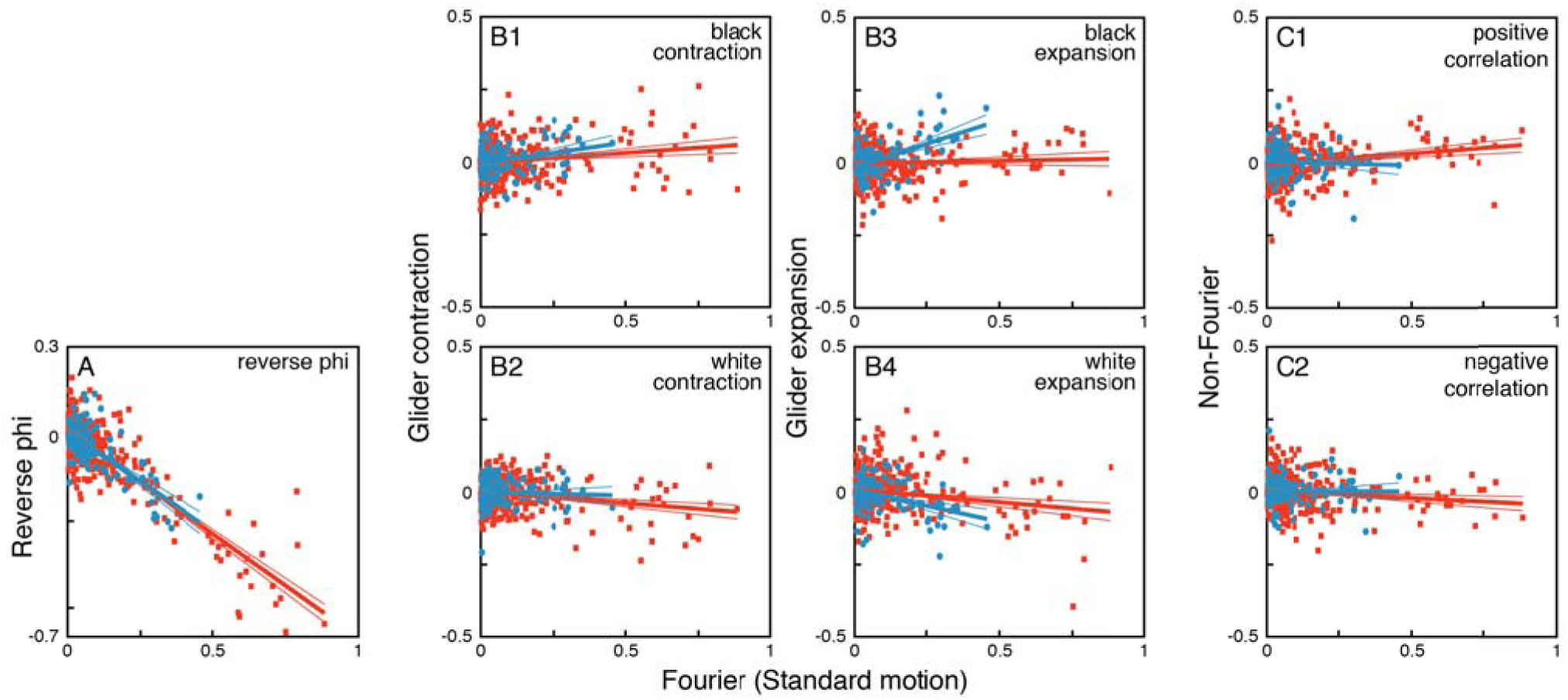
Sensitivities to motion signals are similar at a cellular level. For each unit (total of 53 and 266 in the dragonfly (blue) and macaque (red), respectively), responses to the several motion subtypes were quantified by the direction selectivity index (DI), and plotted against the unit’s DI for Fourier (standard) motion, on the abscissa. Motion subtypes consisted of: (**A**) negatively-correlated Fourier motion (“reverse-phi”), (**B1**) black glider contraction, (**B2**) white glider contraction, (**B3**) black glider expansion, (**B4**) white glider expansion, (**C1**) positively-correlated non-Fourier motion, and (**C2**), negatively-correlated non-Fourier motion. Heavy lines indicate the regression line through the origin; thin lines are 95% confidence limits for slopes (via t-statistics, computed by Matlab’s regstats). Negative values of the DI indicate a direction preference that is opposite to the direction preference for Fourier motion. Lines are draws to the limits of the data along the abscissa.

### Diversity of responses to motion subtypes in simple computational models

The above parallels in response patterns are not merely predictable consequences of imperfect multiplication. Rather, different kinds of deviations from an idealized local motion detector lead to a wide range of motion fingerprints. To demonstrate this range, we carried out simulations of local motion extractors based on a simple scheme of linear spatiotemporal filtering followed by a nonlinearity (Figure 6). In all cases, the linear filter was oriented along a space-time diagonal, and, following the nonlinearity, there was a final “opponent” stage in which matched rightward and leftward signals were compared. Models were presented with the same stimuli used in the experiments, and analysis (see Methods for details) was carried out in a parallel fashion.

**Figure 6.**
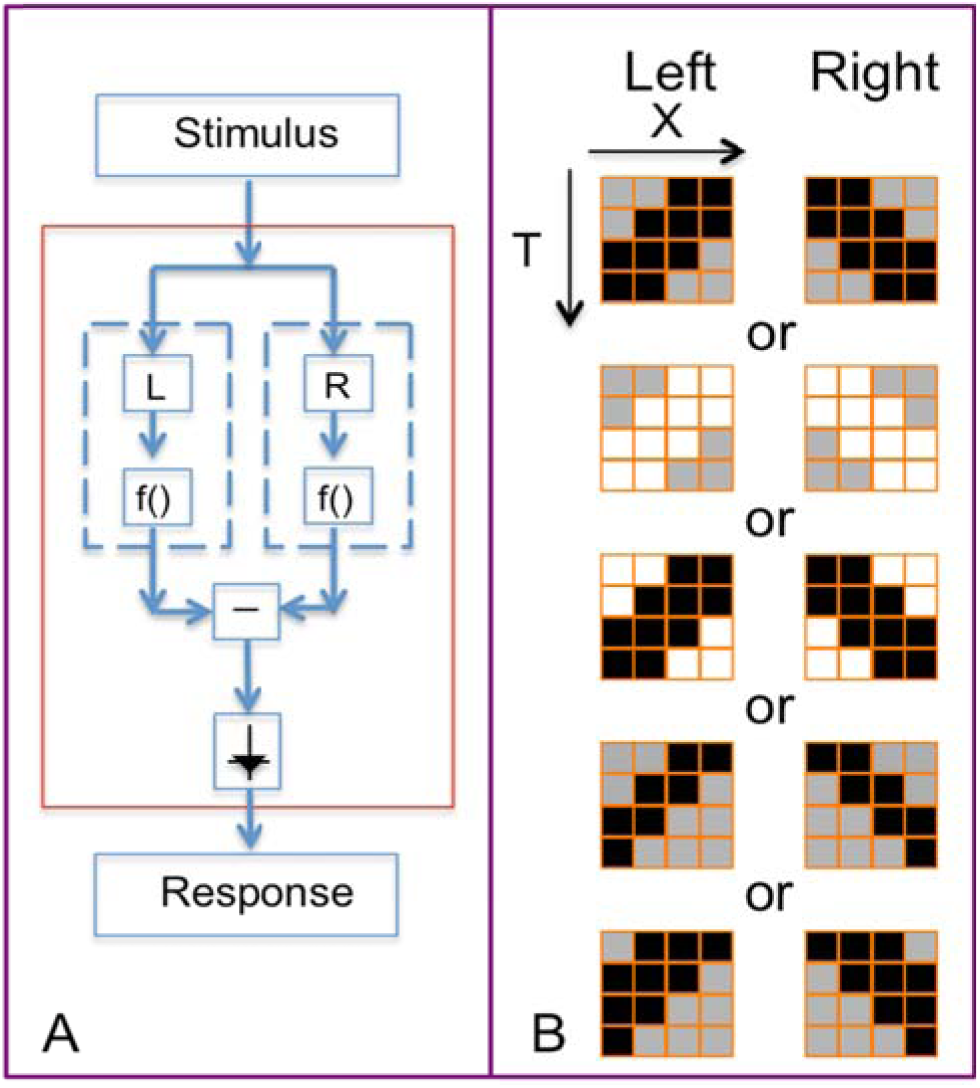
Models for extraction of local motion signals. To explore the diversity of plausible patterns of sensitivity to local motion signals, we constructed simple computational models that generalize the EMD (A). In these models, processing begins with specific linear spatiotemporal filters; several alternative choices are shown in (B). These filters are followed by a static nonlinearity (f in panel A). The model output is the difference between the arm whose filter has a leftward spatiotemporal slant, and the arm whose filter has a rightward spatiotemporal slant. All model filters were constructed on the same spatiotemporal grid as the stimulus set. Colors in the linear filter diagrams (B) indicate the spatiotemporal weights: white: 1, gray: 0 and black: -1.

Results of these simulations are shown in Figure 7, and reveal that the pattern of sensitivity to different motion signal types depends on the nature of the nonlinearity, and, often, on the linear filters. For a strictly quadratic nonlinearity, the model is equivalent the output of a standard EMD, and also to a motion energy model ^9^; as expected, there is no sensitivity to G or NF motion (column 1 of Figure 7). When the nonlinearity deviates from strict squaring, sensitivity to other motion signals emerge. For example, an even-symmetric nonlinearity that deviates from purely quadratic, i.e., *u*^4^ (column 2 of Figure 7), generates responses to NF motion and to G motion. It responds similarly to black and white G motions, but if the linear filter is asymmetric (rows 4 and 5 of Figure 7), it may respond differentially to G contraction and G expansion (but without distinguishing between black and white variants). An asymmetric nonlinearity responds to G motion (columns 3 through 7 of Figure 7) and in most cases to NF motion (columns 3, 5, 6, and 7), and for these nonlinearities, models respond differentially to black and white G motion. For these nonlinearities, the linear filter matters too: dependence on glider polarity is large if the linear filter is unipolar (rows 1, 2, 4, and 5 of Figure 7), and reduced if the linear filter has approximately balanced positive and negative lobes (row 3 of Figure 7). Finally, for linear filters whose spatiotemporal profiles are asymmetric (rows 4 and 5 of Figure 7), responses to expanding and contracting gliders may differ. These differences are more marked when the nonlinearities are asymmetric and accelerating, as seen in columns 4–7 of Figure 7.

**Figure 7.**
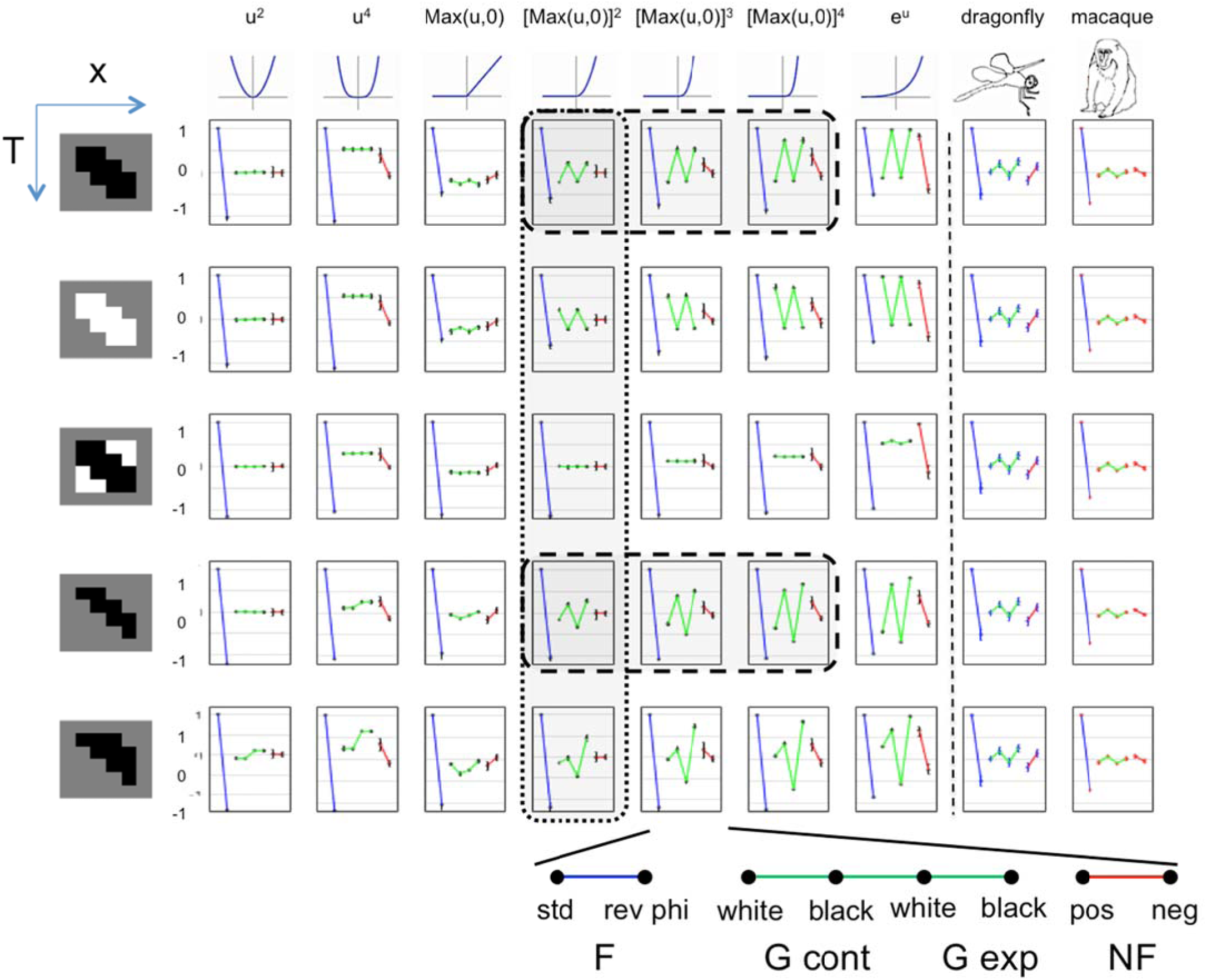
Direction selectivity indices of models for local motion detection show a wide variety of patterns. Each panel shows the direction selectivity indices (DI’s) of a model for local motion detection consisting of a linear spatiotemporal filter, followed by a static nonlinearity, and then an opponent stage (see Figure 6 and Methods for further details). Simulations are carried out for a range of linear filters, shown at the left of each row (colors indicate the spatiotemporal weights: white: 1, gray: 0 and black: -1) and nonlinearities, shown at the top of each column. Model responses show a common pattern for standard Fourier motion (blue), but differ greatly in their pattern of responses to glider stimuli (green) and NF stimuli (red). The response pattern depends both on the choice of linear filter and the nonlinearity. Dl’s (mean and 2 s.e.m.) determined from recordings in dragonfly and macaque are shown in the right-most columns. We show results for nonlinearities that are symmetric or accelerating; results for decelerating nonlinearities (e.g., *e^-u^*) are identical to those for the corresponding accelerating nonlinearities (e.g., *e^u^*) but with the responses to black and white gliders are interchanged. Dashed lines enclose models whose selectivities for the three motion types are consistent with the physiological data. Dotted lines enclose models that use a half-squarer, the nonlinearity that produces responses to glider motion and also motion complexity scores most consistent with the physiologic data (see text and Figures 8 and 9).

Figure S2 presents the simulations of Figure 7 in another format, to facilitate comparison between the different models’ responses to each stimulus type. It emphasizes that models whose responses to F stimuli are identical (upper left panel) have very different responses to the various glider stimuli (four panels on right). Reverse-phi F motion and NF motion distinguish among some models, but the range of model behavior is much larger for the G stimuli.

In sum, simple models of local motion extraction show a wide variety of behavior when probed with a stimulus set that includes the gliders. The common pattern in the neural data – responses to G motion that have the same direction selectivity for black variants, but opposite selectivity for white variants – are seen in only a small subset of these schematic models – those whose linear filters have a predominance of regions activated by negative contrast, and asymmetric nonlinearities (the intersection of columns 4–6 with rows 1 and 4 of Figure 7). However, we stop short of claiming that any one of these models should be taken as “correct”, for several reasons: (i) they are obviously highly stylized, (ii), the agreement with the physiological data is only at a broad qualitative level, without an attempt to account for spatiotemporal tuning or the diversity across neurons, and (iii) the two species differ in the pattern of responses to NF motion (sensitivity to NF in the opposite direction as F for dragonfly, but in the same direction for the macaque) --and both of these patterns can be found in the behavior of these models.

We emphasize that the goal of this modeling was not to fit the neural data, but to show that deviations from an idealized quadratic model can yield many different behaviors, and therefore, that the common pattern of behavior observed across species was not guaranteed *a priori*.

### Opponency in dragonfly, macaque, and models

A striking feature of Figure 7 is that for all models, the DI for reverse-phi motion is negative – that is, the preferred directions for reverse-phi and standard F motion are opposite. This is not surprising, because we built an opponent stage into the model. This reversal in direction selectivity has well-known perceptual correlates in humans: a pattern may appear to move in the opposite direction if its contrast reverses as it moves^22^. This phenomenon is also a well-established feature of the optomotor response in the fruitfly ^3,23^

The prominent opposite-direction response to reverse-phi motion means that a neural response is ambiguous: the same response to motion in one direction may be generated by motion in the opposite direction, if combined with contrast reversal. We hypothesized that a successful motion analysis system needs to resolve this ambiguity. To test this idea, we constructed an index to quantify whether this ambiguity is resolved, and examined how it changed as motion processing progresses, from medulla to lobula in the dragonfly, and from V1 to V2 in the macaque.

The rationale for this index rests on the notion of opponency, as it is not only a pervasive aspect of models of local motion processing but also of visual processing in general^24^. In the EMD^2^, the output – a directional signal – is the result of a two-stage process: a first stage at which spatiotemporal inputs are combined in a nonlinear fashion, followed by an opponent stage that compares two such signals via subtraction. This basic architecture is used in Figure 6, and is also is shared by other extensions of the EMD that extract glider^2^ and non-Fourier motion^15^, and several of the model classes considered by Fitzgerald and Clark^6^. Opponency has a computational benefit, as it is a simple way to eliminate false-positive signals due to uniform flicker and static edges. But this simplicity is the source of the ambiguity mentioned above: after the subtraction of opponent signals, negative correlations in one direction produce the same output as positive correlations in the other. In a purely quadratic motion detector, this ambiguity leads to the reverse-phi phenomenon for standard motion. For opponent detectors with more complex nonlinearities, it will result in black G cues in one direction producing a response similar to that of white G cues in the opposite direction.

Since effective use of motion signals to guide action may require distinguishing between these alternatives, our index (the “motion complexity” (MC) score) is a measure of the deviation from pure opponency, based on comparing direction selectivity for motion subtypes with positive and negative correlations:

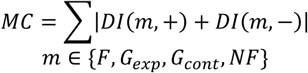
where *DI*(*m,+*) and *DI*(*m,-*) are direction selectivity indices for specific motion subtypes: *DI*(*F,+*) and *DI*(*F,-*) are measured from standard and reverse-phi Fourier motion; *DI*(*G_exp_,+*) and *DI*(*G_exp_,-*) are measured from white and black glider expansion, *DI*(*G_cont_,+*) and *DI*(*G_cont_,-*) are measured from white and black glider contraction, and *DI*(*NF,+*) and *DI*(*NF,-*) are measured from positive and negative-correlation NF motion. Each term in the sum yields a subscore that is 0 only when DI(m,+) and DI(m,-) are equal in magnitude but opposite in sign. The total MC score can only be zero if all of these subscores are 0. This strict opponency means that the nonlinear stage effectively computes a product of contrasts – for example, a pairwise product for a quadratic nonlinearity, and a product of contrasts at multiple points for some of the models considered in ^6^. For other models, at which multiple points interact but not in a purely multiplicative fashion (for example, columns 3–8 of Figure 7, and other models considered by Fitzgerald and Clark^5^), strict opponency will not be present, and the MC score will be larger than 0.

Figure 8 shows the MC score distributions of neurons recorded in the two species. Consistent with our hypothesis, the distribution of MC scores shifts towards higher values in the second processing area (lobula or V2), compared to the earlier processing (medulla or V1). This shift is present for each of the four MC subscores (Figure 5A), and is significant (p<0.001 for V1/V2, two-tailed Wilcoxon ranksum test) for the total MC score in the macaque (Figure 8B).

**Figure 8.**
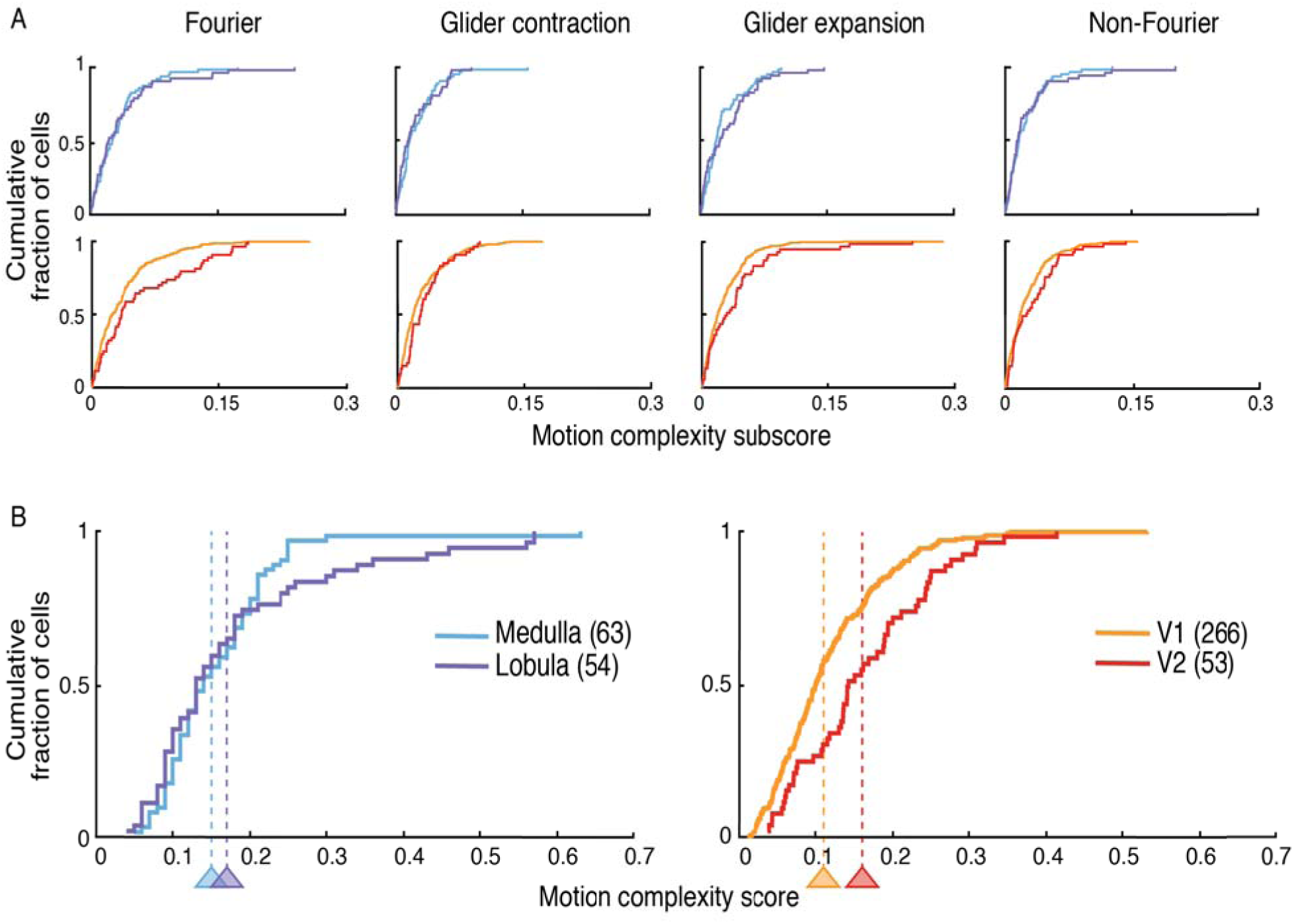
In dragonflies and macaques, the ambiguity of opponency is partially resolved as motion processing unfolds. The motion complexity score (MC, see text) indicates whether an individual neuron's response can resolve the ambiguities inherent in an opponent calculation (see text): neurons with higher MC scores are able to distinguish a positive motion signal in one direction from a negative motion signal in the opposite direction. We determined the distribution of MC subscores (**A**) and the total MC score (**B**). In dragonflies (left) and macaques (right), the distribution of MC scores shifts towards higher values from medulla (light blue) to lobula (dark blue) and from V1 (orange) to V2 (brown). Thus, as motion processing unfolds, neural computations progress beyond opponency, allowing neurons to distinguish positive spatiotemporal correlation in one direction from negative spatiotemporal correlation in the opposite direction. In B, vertical dashed lines indicate mean MC score.

The MC score is another way in which models may manifest diverse behavior. Figure 9 demonstrates this for the models considered above, and compares these model MC scores with those obtained in dragonfly and macaque. MC scores in the range of the experimental data are found for two kinds of nonlinearities: the full squarer (column 1 of Figure 7), which does not produce glider responses, and the half-squarer, max(0,x)^2^, which does (column 4 of Figure 7). Thus, the overall characteristics of the physiologic results -- the pattern of glider sensitivity with a low MC score – are accounted for by only a small fraction of model variations.

**Figure 9.**
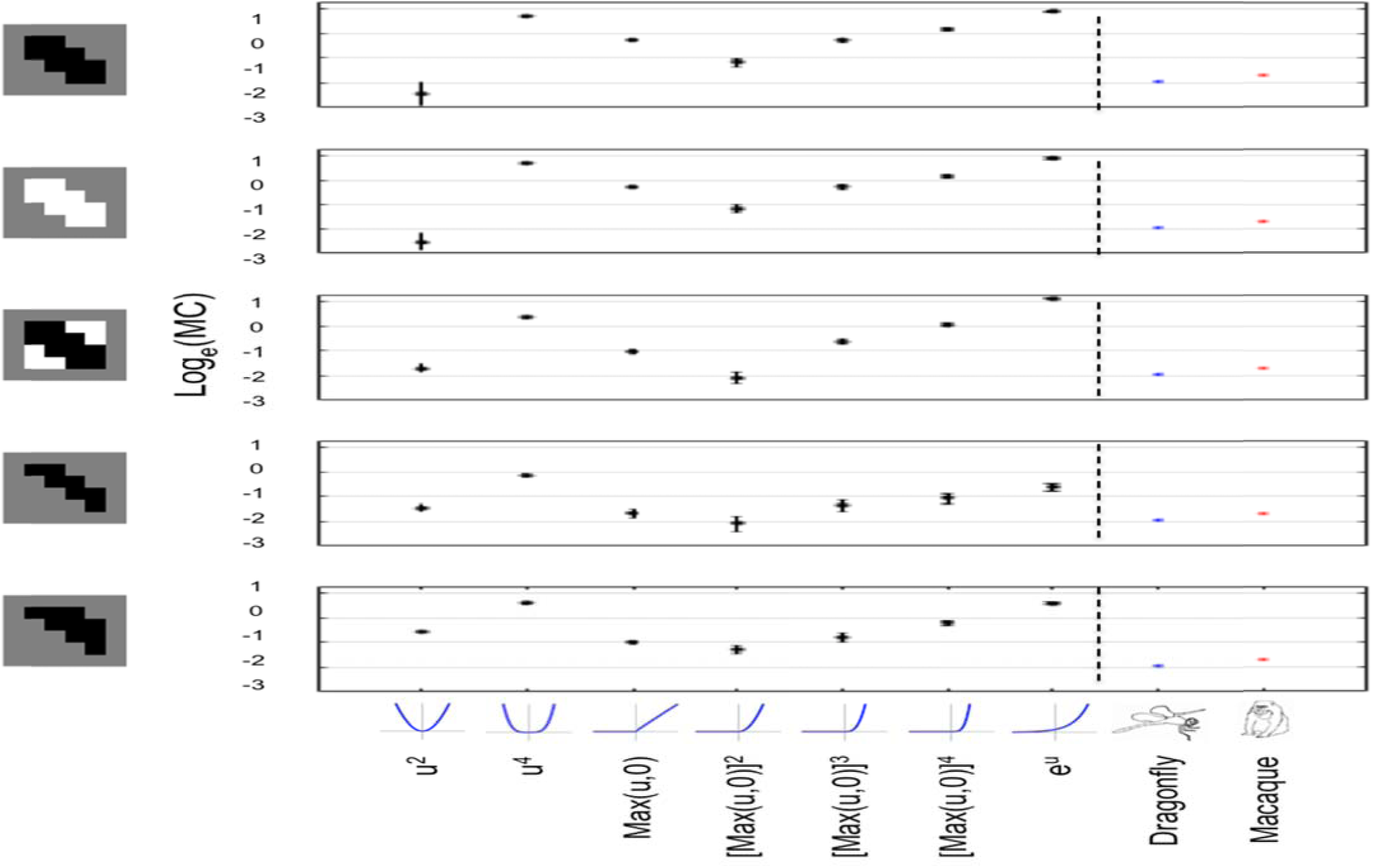
Motion complexity scores of models for local motion span a wide range. MC scores (median and 2 s.e.m.) are calculated for the models of Figure 7, and compared with MC scores determined from the physiological recordings. MC score depends primarily on the model’s nonlinearity, and many nonlinearities lead to MC scores above the range of the recordings. MC scores for decelerating nonlinearities (e.g., *e^-u^*) are the same as for the corresponding accelerating nonlinearities (e.g., *e^u^*), and are not shown. Note that the ordinate is logarithmic, and in some cases the error bars are not visible because they are smaller than the symbols.

## Discussion

The seemingly effortless nature of visual motion analysis gives little hint of the complexity of neural computations that may be required to support them.

There are two basic reasons for this complexity: the diversity of the behavioral repertoires that rely on motion cues, and the diversity of the motion cues themselves. The diversity of the behavioral repertoire is clearly evident in predator species. Predator species – along with many others – use motion signals to help control locomotion or flight, which requires extracting a global flow field; for this purpose, point-to-point variations in local motion signals are effectively a source of noise to be averaged out. This use of motion has been the focus of a large body of work, especially in insects ^4,25^ But predator species, including predator insects, also need to identify and pursue prey. For these behaviors, and for the use of motion to recognize objects in general, point-to-point variations in local motion signals are the critical signals, and the overall flow field, which may be generated by self-motion, can be ignored as noise.

With regard to the motion cues themselves, diversity arises because of the two basic ways that local motion signals can be generated: self-motion, and motion of an object with respect to its background. These processes differ in their statistical characteristics, and their net effect on the retinal image is complicated by the geometry of occlusion and 3D projection. Conversely, although all visual motion signals entail a change in the visual image across time, not every spatiotemporal change stems from motion. For example, flicker typically stems from a change in the illuminant or shadowing, and needs to be separated from spatiotemporal changes that originate from motion.

Despite this potential for complexity, a simple model, the Hassenstein-Reichardt model, also known as the elementary motion detector (EMD), has been very successful in providing a basic understanding of motion analysis^2,4,26,27^. The first stage of the EMD computes a two-point spatiotemporal correlation: it multiplies the visual signal at one point (say, A) by the visual signal at a nearby point (say B), at a slightly later time. If there is motion from A to B over this time interval, then these two signals values will be strongly correlated.

To eliminate spurious signals that are not due to motion, the EMD has a second stage, which uses an “opponent” strategy: it subtracts the correlation computed in the A-to-B direction from the correlation computed in the B-to-A direction. If the source of the correlation is motion from A to B, then the correlation in the B-to-A direction will be weaker, and will not result in cancellation. But if the source is diffuse flicker, the correlations will cancel, and no net signal will be produced. In addition to accounting for a great deal of behavioral data^2,22,26,27^, the EMD model has the attractive feature that it maps directly onto the lamellar anatomy of the insect visual system. Thus, the opponent strategy is an elegant computational solution to disambiguating a simple kind of directional motion signal from confounding flicker^4,28^ – though it has the side effect of confounding true motion in one direction with reverse-phi motion in the opposite direction.

As noted, the first stage of the EMD consists of a multiplicative interaction between two signed quantities, the local contrasts at A and B. However, signed (4-quadrant) multiplication is not a natural operation for biological components. In flies, the circuitry that executes the interactions postulated by the EMD only carries out approximations to multiplication^8,28,29^. Individual neural interactions are dedicated to a single polarity of stimuli, and the interactions for positive and negative-polarity stimuli are not symmetric^30^. That is, the interaction between signals in a biological EMD is only approximately multiplicative.

This leads to two possibilities: (i) an approximate multiplication may suffice for local motion analysis, and the evolutionary, developmental, or metabolic costs of circuitry required to make it more precise exceed the computational advantages of strict multiplication, or (ii) the deviations from multiplication are in fact advantageous, and point to a biological algorithm for motion analysis that is better-tuned to visual tasks than the standard EMD.

Several lines of evidence now indicate that the latter is the case. First, it has long been recognized that there are cues to motion are undetected by the computation specified by the EMD: these include four-point correlations (“non-Fourier” motion, ^13^), and, more recently, three-point correlations (“glider motion”, ^11^). These cues are consistently present in the natural environment^12^, and arise as a result of black-white asymmetry^5,30^, occlusion, non-rigid motion, and motion in 3D. Second, these cues are non-redundant with the cues extracted by the EMD, so there is an advantage to exploiting them^5,30^. Third, a wide variety of species show behavioral responses to these cues (*Drosophila:* ^3^ zebrafish: ^14^; human: ^11,15^). Fourth, when challenged with moving naturalistic 1D patterns, specific deviations from the EMD that are sensitive to glider motion lead to more veridical estimates of optic flow, and are better predictors of optomotor behavior^6^ in the fruitfly. Here we show that the parallel deviations from the EMD are present in two visual specialists that use vision to acquire and intercept objects – the dragonfly and the macaque. These parallel deviations primarily relate to responses to glider motion, and consist of a similarity in the proportions of neurons that respond to different motion signal subtypes (Figure 4), and the way that these signals are combined within individual cells (Figure 5). Additionally, both dragonfly and macaque demonstrate a shift away from pure opponency as processing unfolds (Figure 8). However, there are two identifiable differences between the species: in the dragonfly, responses to black glider expansion are larger and more strongly correlated with standard two-point motion (Figure 5B3), and responses to non-Fourier motion tend to have an opposite relationship to the response to standard motion in the two species (Figure 5C1, 5C2).

It is notable that the parallels we found are present at the level of circuitry (i.e., the way that individual cells respond to different kinds of motion signals), and not just at the level of the overall computation performed. The latter might be expected based on normative considerations^5,31^, but the former is rather remarkable, given these species’ overwhelming differences in brain size, neuron number, and organizational plan.

Along with these obvious anatomical differences between species, there are differences in the framework in which local motion computation takes place – and these differences amplify the implications of the parallels identified here. In dragonfly^10^, as in Drosophila^8^, directional selectivity arises in the retina. Consistent with the EMD model, this computation involves a nonlinear interaction of a pair of luminance signals separated in space and time. An analogous nonlinear interaction of local luminance signals is also present in mammalian retina^31,32^, resulting in directionally-selective retinal ganglion cells. In both cases, these local interactions are effectively modeled as a multiplication, and the challenge of implementing a signed multiplication in neural circuitry is met in both cases by a separation of signals into ON and OFF channels ^3,30,31,33,34^

While primates have directionally-selective retinal neurons^35^, these are believed to play a minor role in high-resolution visual tasks: in the primate, most direction selectivity appears to arise in the cortex^10^. Cortical direction selectivity is thought to arise by comparing two “motion energies”^9,36^. In the idealized form of the computation, each motion energy is the square of the output of a spatiotemporal filter, and these opposing motion energies are subtracted. At first glance, subtraction of motion energies seems rather different than multiplication – but because of the algebraic identity (X+Y)^2^-(X-Y)^2^=4XY, it leads to the same result ^9^.

Thus, in the primate, the Reichardtian need to implement signed multiplication is circumvented, but it is replaced by a need to compute squares, and to have closely matched neural filters. Just as the presence of responses to G motion implies that multiplication in the insect retina is not precise, these responses imply that the motion energy computation in the primate is also approximate. The deviations of the neural computations from their simplified ideals arise in different ways, but nevertheless lead to the same pattern of G sensitivity: neurons with directionally-selective responses to standard motion tend to prefer black G expansion and contraction in the same direction, and white G expansion and contraction in the opposite direction.

Finally, it is tempting to speculate about the functional benefits of this pattern of G responses. If we assume that objects tend to be darker than their backgrounds (as is typically the case when the dominant background is the sky), then objects moving in depth will generate a black G signal that is informative about their motion. White G signals, on the other hand, would arise from changing sizes of gaps – e.g., spaces between branches – rather than from motion of objects – and thus, should not be used to reinforce an F motion signal. Recognizing that this viewpoint is both speculative and highly simplified, it’s interesting to note that black G expansion signals are somewhat stronger in the dragonfly than in the macaque (Figure 5B3), perhaps owing to their airborne lifestyle and greater need to intercept dark objects moving against a lighter background.

In sum, we compared the analysis of visual motion at the neuronal level in the central nervous systems of two extremely different visual specialists, whose evolutionary and anatomical divergences are profound: the dragonfly and the macaque monkey. We conclude that ethological demands drive biologic motion processing in these species to a convergent solution at the neuronal level despite major differences in their phylogenies and the architectures of their eyes and brains. This convergence provides strong evidence that neuronal implementation of local motion computations are finely tuned to the statistics of the visual environment.

## Materials and Methods

### Physiology and Recording Methods

#### Dragonfly

##### Physiological preparation

Multiple dragonfly genera were used in these experiments including *Anax junius*, *Aeshna verticalis* and others, which came from one of two sources: wild-caught (Ithaca, NY; May-October 2013) and laboratory reared (Carolina Biological Supply Co.). Recordings were made from a total of 26 animals.

After capture, wild-caught dragonflies were held for short periods that did not exceed 15 hours in the laboratory before use. Laboratory reared animals arrived as penultimate nymphs and were raised in house (12:12 light/dark cycle; 80% humidity; 27 deg C) in individual containers and fed a diet of mosquito larva until eclosion. All experiments with laboratory reared dragonflies took place no more than 48 hours after eclosion.

Just prior to the start of each experiment, dragonflies were cold anesthetized for 2–4 minutes in a freezer (-4 deg C). Dragonflies were then restrained and affixed to a plastic post using Kerr dental sticky wax (58 deg C melting point, Syborn Kerr, Emeryville, CA, USA) heated by a cool soldering iron (Antex model C, Antex (Electronics) Limited, Travistock, Devon, UK) with the voltage limited to 55V using a variable transformer (Powerstat type 3PN116B, The Superior Electric Co., Bristol, CT, USA). The animal was positioned ventral-side-down on the post, and the head was tilted downward such that the dorsal high-acuity fovea was pointed towards the screen. A small flap of cuticle on the anterior portion of the head, between the eyes and the neck (thorax), was removed to expose the right optic lobe of the brain over the medulla and lobula. A drop of fresh extracellular saline solution containing (in mM): 185 NaCl, 4 KCl, 6 CaCl2, 2 MgCl2, 10 HEPES, and 35 D-glucose (solution adjusted to pH 7.2 with NaOH and 430 mOsm with glucose; ^37^) was placed on the brain at least every 30 minutes.

##### Recording and visual stimulation

Recordings were made using tungsten microelectrodes (4MΩ; MicroProbe Inc., Gaithersburg, MD, USA) mounted to stereotactic micromanipulators (Narishige International USA, Inc., East Meadow, NY, USA) and advanced using a hydraulic microdrive (Model 607W, David Kopf Instruments, Tujunga, CA, USA) at 1μm steps once inserted into the brain. Electrode placement into the medulla or lobula was determined visually by the experimenters using anatomical landmarks ^38^. Electrical activity was acquired via an extracellular headstage (Model 1800 A-M Systems, Sequim, WA, USA) and amplified 10,000x and filtered (100Hz-5000Hz bandpass, 60Hz notch) using a differential AC microelectrode amplifier (Model 1800 A-M Systems, Sequim, WA, USA), followed by an A/D converter (NI PCI-MIO-16E-1, National Instruments, National Instruments, Austin, TX, USA) fitted with a breakout box (NI BNC-2090, National Instruments, Austin, TX, USA). All recordings were made at 15kHz sample rate using the Spike Hound data acquisition software (formerly called g-Prime; ^39^ on a computer running Windows 7 (64-bit; Microsoft Corporation, Redmond, WA, USA). All recordings were done on an air table (Micro-G, Technical Manufacturing Corporation, Woburn, MA, USA) with a custom-built wire-mesh Faraday cage.

Visual stimuli were presented using a conventional 37 by 22cm LCD computer monitor (ViewPanel VE150m, ViewSonic, Walnut, CA, USA) at a refresh rate of 60Hz and resolution of 1920x960 pixels and mean luminance of 53 cd/m^2^. Animals were positioned 22.8 cm from the screen, which resulted in stimulus check sizes that were approximately 2.5 degrees. Note that stimulus parameters were not optimized to the tuning of the recorded neurons, due to the limited stability of the extracellular recordings (typically about 30 min). Stimuli were presented using a custom-made video player. The program presents the stimuli and synchronizes the recordings.

Once acquired, single units were isolated using a customized version of WaveClus ^40^. WaveClus processing entailed a bandpass filter (300–6000 Hz), thresholding to detect candidate spikes, decomposition of each candidate spike into eight Haar wavelet features, and clustering of spike events based on the wavelet coefficients for each. For each recording, sorting was carried out using different amplitude thresholds and cluster partitioning until we were confident that single units were isolated using similar criteria as in the macaque.

#### Macaque

##### Physiological preparation

Standard acute preparation techniques were used for electrophysiological recordings from V1 and V2 of cynomolgus monkeys (*Macaca fascicularis*) weighing 2.2 to 10 kg (12 males, 1 female). All procedures were approved by the Animal Care and Use Committee of the Weill Cornell Medical College and consistent with Institutional and National Institutes of Health guidelines for the care and experimental use of animals. Procedures were previously described in detail ^20,41–43^ and are summarized here. Animals were premedicated with atropine (0.05 mg/kg, i.m.; Henry Schein). Following ketamine (Ketaset, 10 mg/kg, i.m.; Fort Dodge Animal Health) or Telazol (4 mg/kg, i.m.; Fort Dodge Animal Health) and under isoflurane (1–2%; Hospira) surgical anesthesia, an endotracheal tube was placed, catheters were inserted in both femoral veins and one femoral artery, and a craniotomy was made near coordinates P10, L15. During recording, anesthesia was maintained with propofol (PropoFlo, 2–20 mg/kg/h, i.v.; Abbott) and sufentanil (Sufenta, 0.1–1 micrograms/kg/h, i.v.; Janssen) and neuromuscular blockade was established (following all surgical procedures) with vecuronium bromide (0.25 mg/kg, i.v. bolus, 0.25 mg/kg /h, i.v.; Bedford Laboratories) or rocuronium bromide (1.5 mg/kg, i.v. bolus, 1–1.5 mg/kg/h, i.v.; Mylan Institutional). During the experiment, heart rate and rhythm, arterial blood pressure, body temperature, end-expiratory pCO_2_, urine output, and EEG were monitored. Routine maintenance included intravenous fluids, periodic O_2_ supplementation, antibiotics, dexamethasone, application of local anesthetics to surgical sites, ocular instillation of atropine (1%; Bausch & Lomb), and flurbiprofen (Ocufen, 0.03%; Allergan), and periodic cleaning of the gas-permeable contact lenses (Metro Optics) behind 2-mm artificial pupils. Lenses with spherical correction, subsequently adjusted to maximize the responses of isolated single units to high-spatial-frequency visual stimuli, were used to focus the stimulus on the retina. With these measures, the preparation remained physiologically stable for 4–5 days.

##### Recording and visual stimulation

Through a small durotomy over V1 and/or V2, an array of 3 or 6 tetrodes (quartz-coated platinum-tungsten fibers; Thomas Recording, Giessen, Germany) was inserted, avoiding surface blood vessels via a custom headstage (Thomas Recording) that allowed for adjustments of the array geometry and independent lowering of each tetrode. Signals from each tetrode channel were amplified, filtered (0.3–6 kHz), and digitized (25 or 30.303 kHz). Once spiking activity from one or more units was encountered, the region of the receptive field(s) was hand-mapped and then centered on the display of a 21-inch gamma-corrected CRT monitor, (1280 × 1024 raster, 100 Hz refresh), either a ViewSonic G225f 21-inch monitor (mean luminance 47 cd/m^2^) or a Sun GDM5410 21-inch monitor (mean luminance 46 cd/m^2^) at a distance of 114 cm. Control signals for the monitors were provided by a PC-hosted system optimized for OpenGL (NVidia GeForce3 chipset) programmed in Delphi.

Following hand-mapping, computer-controlled presentation of drifting sinewave gratings were used to characterize neural responses, including orientation tuning, spatial frequency tuning, temporal frequency tuning, and the contrast-response function. One unit whose extracellularly-recorded action potential was identifiable by on-line spike sorting was chosen as the “target neuron.” This neuron’s orientation and direction preference was used to determine the orientation of the motion stimuli (see SM §2), and its spatial frequency optimum was used to determine the check size (approximately 2 checks per lobe of the optimal grating). For the motion stimuli, contrast was always 1.0 and stimulus velocity was always 10 checks per second. With the typical check size of 0.2 deg (rarely less than 0.1 deg or greater than 0.5 deg), stimulus velocity was 2 deg/sec.

Offline, recordings were spike-sorted as described in ^41^, based on automated clustering via KlusterKwik ^44^ operating on 17 features (peaks and troughs on each of the four channels, the first eight principal components of the wave shapes, and spike time), followed by hand merging and reclustering in Klusters. Criteria for single neuron isolation included waveform shape, its gradual change over time, and the number of refractory period violations.

##### Histology

Procedures were identical to that of ^41^. In brief, lesions were made after all recordings were completed, and, following a waiting period of 1 h, the animal was deeply anesthetized and perfused (4% paraformaldehyde; EMS). The border between V1 and V2 was identified via the distinct appearance of layer 4 in V1 and its disappearance in V2. We marked each unit as certain V1, certain V2 or uncertain V1/V2. The latter units were only included in the analyses that were not subdivided according to area.

### Motion Stimuli

For both species, motion stimuli consisted of a temporal sequence of “motion blocks.” Each motion block consisted of a segment containing a particular motion signal subtype in one direction (1500 ms duration, containing 15 frames of 100 ms each, followed by 500 ms of gray (50%)), followed by a similar segment containing the same motion signal subtype in the opposite direction. The opposite-direction segments paired in each motion block were presented in pseudorandom order. We analyzed responses to 8 kinds of motion blocks that contain Fourier, glider, and non-Fourier signals (Figure 1), along with 5 other kinds of motion blocks that served as controls (These control motion blocks are based on three- and four-point gliders that did not yield strong percepts of motion ^11^, and did not yield direction-selective neural responses). For each kind of motion block, movies were generated with pseudorandom seeds and contained, on average, 50% black checks and 50% white checks, according to the procedure of Hu and Victor ^11^. These motion blocks were each repeated 32 times, totaling 26 minutes. In the macaque, this sequence was repeated up to four times. Each segment in each motion block was constructed with a different random seed.

Example movies of all motion stimuli can be found in the video supplement.

Note that all stimuli (Fourier, Glider, non-Fourier, and the controls) were defined by “templates” (spatiotemporal correlations) confined to a 2x2x2 spatiotemporal volume of checks (Figure 1), so that they had comparable spatial and temporal extent. As a consequence, the non-Fourier motion stimuli used here consist of motion of an edge that is parallel to the motion direction. Other studies of non-Fourier motion in mammalian V1 and V2 ^16,17^ used non-Fourier stimuli defined by contours that were orthogonal to the motion direction, and this may account for the higher fraction of neurons that were found in those studies to be sensitive to non-Fourier motion.

As mentioned in the main text, the level of sensitivity to Fourier motion is different across species, with lower levels seen in dragonfly neurons than in the macaque. This difference may be due to the fact that for the macaque, stimuli were optimized for velocity (by adjusting spatial frequency), as well as for preferred orientation and contrast, but no optimization was done for the dragonfly (see Methods §1.1 and §1.2 above). Another possible factor is that dragonflies are known to capture changes in moving images at more than 200Hz rate ^45^, but stimuli were only updated at 10 Hz in both species (display frame rate: 60 Hz for dragonfly, 100Hz for macaque). The reason for this is that the strategy for isolating motion types (Figure 1) requires stepwise updates (i.e., a discretization of stimuli in time), and we chose this discretization to be identical in the two species.

### Data analysis

To determine whether a neuron had a directionally-selective response for each motion subtype (e.g., white glider expansion, and other examples in Figures 2 and 3), we compared the total number of spikes that occurred between 50ms and 1600ms following the onset of the motion movie (duration, 1000 ms) in the two directions within each motion block (paired t-test, criterion p=0.05).

A neuron was considered to have a directionally-selective response to a kind of motion (Figure 4) if it had a directionally-selective response to any of its subtypes: for Fourier motion, the subtypes consist of standard and reverse-phi motion; for Glider, the subtypes consist of expansion and contraction, each black or white; for non-Fourier motion, the subtypes consist of positive and negative correlation.

For each motion subtype, direction selectivity was quantified by a direction selectivity index (DI), given by equation 1: *DI=*(*M_pref_-M_non-pref_*)*/* (*M_pref_* + *M_non-pref_*). Directions were labeled “preferred” and “non-preferred” based on their responses to Fourier (standard) motion. Thus, for a neuron that responded to another subtype with the opposite direction preference compared to its response to Fourier motion, the DI was negative.

All neurons with a firing rate of 1 Hz or more were analyzed.

### Simulations

To understand the pattern of sensitivities to motion signals that would arise from local motion detectors that are not strictly multiplicative, we simulated responses of simple models to the same stimuli used in the experiments. Models (Figure 6) consisted of two opponent arms: one containing with a linear filter that is oriented along a space-time diagonal to the left, and a mirror-symmetric one that is oriented along a space-time diagonal to the right. For simplicity, filters were constructed on the same grid as the stimuli, and filter weights were all +1, 0, or -1. Convolution of the stimulus with these filters was computed in discrete time. The results of the convolution were the inputs to identical static nonlinearities, and the outputs of the nonlinearities were subtracted to determine the model response. As shown in Figure 6, we considered filter profiles that had positive, negative, and mixed lobes, and nonlinearities that deviated from squaring in several ways (e.g., symmetric vs. asymmetric, rectifying vs. smoothly accelerating).

To compare model responses with neural responses, we placed the model in a random location on the stimulus array, and computed its response to the same set of 32 sequences (10 frames in each direction) used in the experiment. Responses to stimuli moving in opposite directions were compared to determine the direction selectivity index (DI) and the motion complexity score (MC). Since each stimulus sequence was finite, these quantities depended on the placement of the model on the stimulus. This variability is reflected in the error bars (2 s.e.m., estimated from 100 random placements) shown in the Results (Figures 7 and 9).

## Author Contributions and Notes

E.I.N., R.R.H., and J.D.V. conceived and planned the experiments. E.I.N., G.M., P.S.S., Q.H. and J.D.V. carried out the experiments. E.I.N., and J.D.V. planned and carried out the simulations. E.I.N., G.M., P.S.S, J.R.G., R.R.H., and J.D.V. analyzed the data and contributed to the interpretation of the results. J.D.V. and R.R.H. took the lead in writing the manuscript. All authors provided critical feedback and helped shape the research, analysis and manuscript.

## Acknowledgments

We are very grateful to Cole Gilbert, Daniel Zurek, Robert Wyttenbach, Melissa Warden and Jose Manuel-Alonso for their insights and comments on earlier drafts. EIN was supported by Cornell University's Tri-Institutional Training Program in Computational Biology and Medicine. This work was supported in part by NIH EY9314 and EY7977 to JDV and by NIH 5R01DC 10338 to GM, PSS and RRH

## Supporting Information

### S1. Appendix. Additional Analyses – Figure Supplement

#### S1.1 Alternative criterion for sensitivity

In Figure 4 of the main text, we defined a neuron to be sensitive to a given kind of motion if it was had a significant (p<0.05) directionally selective response to any of its subtypes (Figure 1). Here, we include a Bonferroni correction to account for the multiple subtypes: the significance cutoff for Fourier motion (two subtypes) is p<0.025, for glider motion (four subtypes) the cutoff is p<0.0125, and for non-Fourier motion (two subtypes) the cutoff is p<0.025. Note that this correction is conservative, since it assumes that responses to motion signals within a type are uncorrelated. As in the primary analysis (Figure 4), the pattern of motion sensitivity is very similar between dragonfly and macaque.

**S1.**
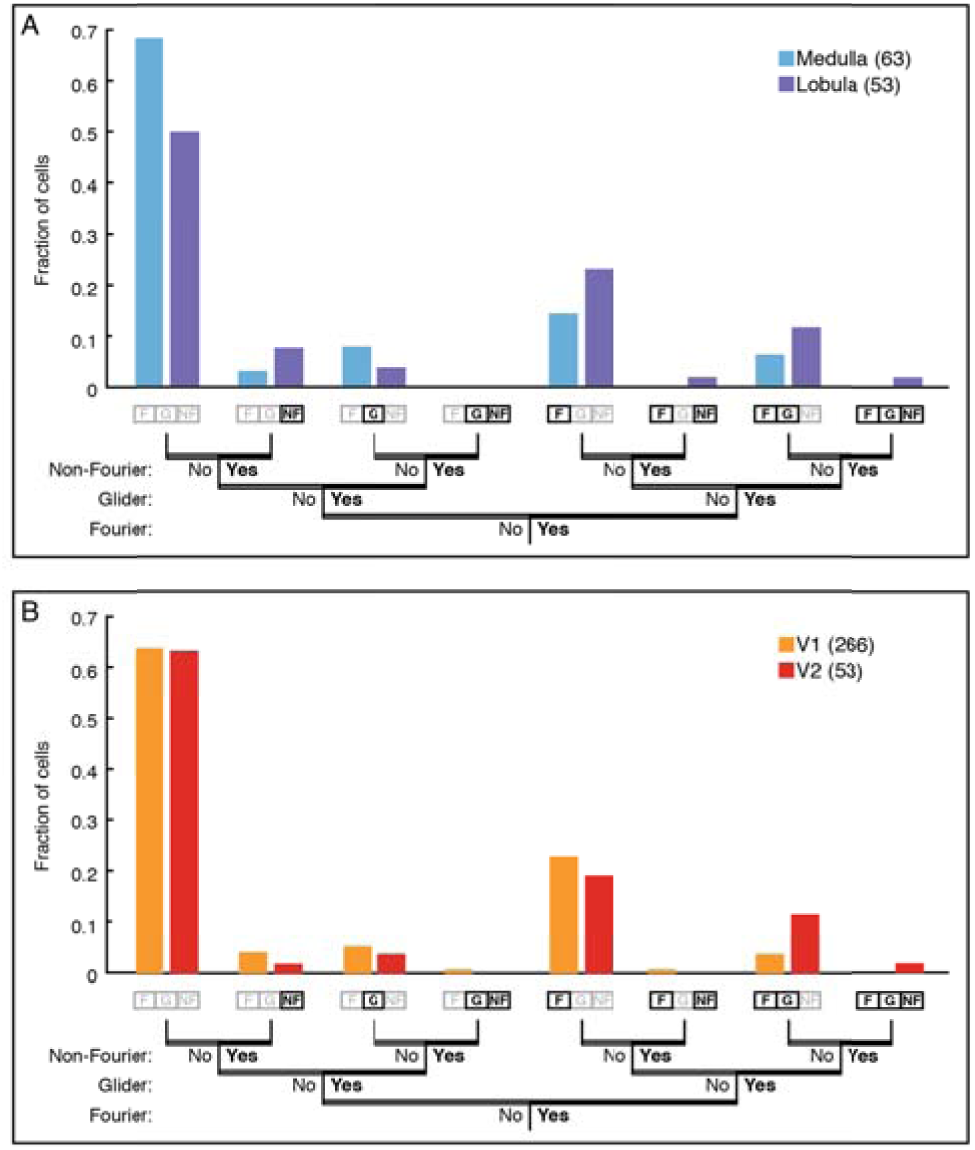

#### S1.2 Alternative view of selectivity to local motion types

In Figure 7 of the main text, we presented the sensitivity of a range of models to F, NF, and G motion types, showing that the motion fingerprint depended on an interaction between the linear filter and the nonlinearity. Here we present an alternative view of the simulation results, facilitating a comparison of how the different motion subtypes distinguish among models.

**S2 Figure.**
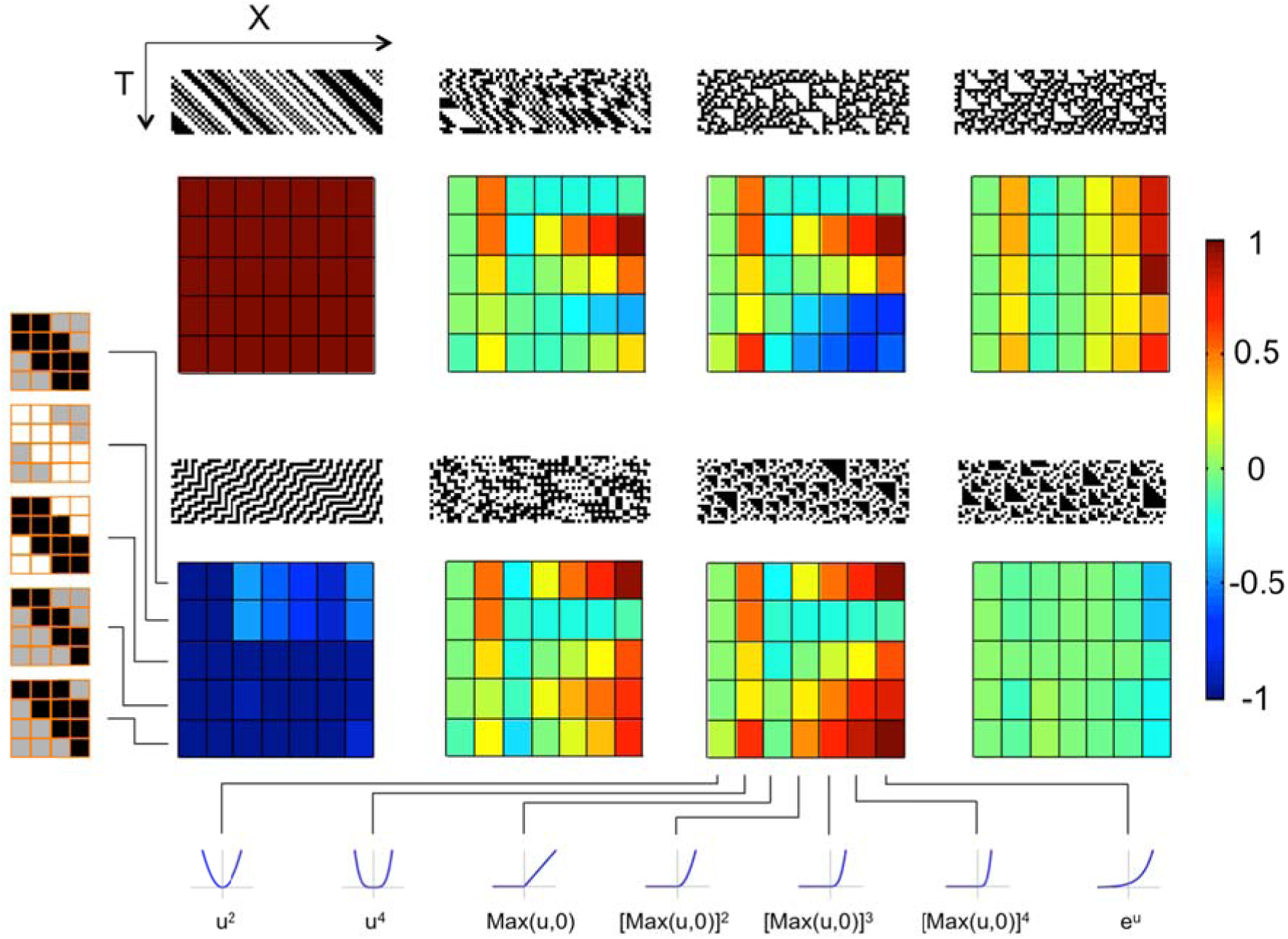
F, G, and NF motion subtypes have varying abilities to distinguish among models. For each motion subtype (top row: positive correlation; bottom row: negative correlation), we show the direction selectivity for a range of models as a colormap under the XT diagram for each motion type. These models consist of the 5 linear filters combined with the 7 nonlinearities considered in Figure 7 of the main text. Direction selectivity indices are normalized to 1 for F motion (top row, left). For reverse phi (bottom row, left), all models have a negative direction selectivity. For NF motion (right column), model responses are somewhat more varied, but depend primarily on the shape of the nonlinearity. For G motions (two middle columns), direction selectivity depends on an interaction between the linear filter and the nonlinearity, and can be either positive or negative. Compared to the diversity of responses to reverse-phi and NF motion types, the diversity of the responses to G motion provides the greatest possibility of distinguishing among models. Filters displayed as in Figure 7 of the main text.

### S2 Appendix. Video supplement

The video consists of short clips of stimuli used in these experiments. In each case motion is first to the right, then to the left. The final clip is a random checkerboard.

1. Fourier standard motion
2. Fourier reverse phi
3. Glider black expansion
4. Glider black contraction
5. Glider white expansion
6. Glider white contraction
7. Non-Fourier positive correlation
8. Non-Fourier negative correlation
9. Random

